# Genomic and oceanographic evidence of reef connectivity across the western and central Indian Ocean

**DOI:** 10.64898/2026.10.22.761076

**Authors:** Catherine E.I. Head, Luigi Colin, Phil Hosegood, Michelle L. Taylor, Maria Belen Arias-Mella, April Burt, Anna Koester, Margaux Steyaert, Tom Bridge, Chris Yesson

## Abstract

Connectivity among coral populations, via larval dispersal, is one of the key mechanisms supporting reef recovery after climate-driven bleaching events. This study examines reef connectivity of remote atolls of the understudied central and western Indian Ocean seascape, focusing on a key reef-building species, *Acropora aff. tenuis*. Using genomic data from ultra-conserved elements alongside particle-tracking simulations, we investigate patterns of population structure and gene flow, and the influence of ocean currents. Results show asymmetrical connectivity: strong gene flow occurs within the Chagos Archipelago and between Chagos and the Inner Seychelles, highlighting their importance as regional larval sources. In contrast, the Outer Seychelles mainly functions as a larval sink, and Îles Glorieuses appears to be the least connected. Whilst population structure suggests a partially permeable dispersal barrier between the Inner and Outer Seychelles. Differences between genetic patterns and simulated ocean-driven dispersal suggest that ocean currents are not the only important driver of connectivity patterns. Additional factors, such as stepping-stone dispersal among atolls, fine-scale oceanographic processes, and species biology likely play important roles. Overall, we find widespread genomic connectivity across this seascape, which has positive implications for reef resilience, though future resilience will depend on frequency and severity of future bleaching events.

## Introduction

Global climate-change induced increases in sea surface temperatures (SSTs) are now the biggest single threat to coral reefs (1,2). Reefs in the Indian Ocean (IO) suffered severe mortality from the third global coral bleaching event in 2014-2017 (3,4), with even protected areas isolated from local anthropogenic pressures such as Aldabra Atoll and the Chagos Archipelago experiencing severe bleaching and mortality (5–8). While still recovering from this event, another marine heatwave affected coral reefs throughout the IO in early 2024 (Cordio, 2024), as part of the fourth global coral bleaching event, the full impacts of which are not yet documented. This marine heatwave in early 2024 was driven by El Nino and positive Indian Ocean Dipole (IOD) phases. Positive IOD phases result in warmer than normal SSTs in the Western Indian Ocean (WIO), altering the direction of the atmospheric circulation, with associated biological impacts (9)

The outlook for reefs globally is disheartening, with global climate models predicting that >75% of reefs will experience annual severe bleaching before 2070 even under the most optimistic climate change scenario (RCP4.5) (10,11). However, it is difficult to incorporate adaptive potential of corals into ecological models, and recent studies suggest that natural selection could allow coral populations to persist if global warming is kept to ∼2°C (12). Ultimately global climate change must be tackled to bring emissions down and give reefs the best chance of survival (1,13,14). Meanwhile, local management efforts can build resilience and restoration techniques can help re-build damaged reefs, but with limited resources these efforts must be directed to be as effective as possible (15–17).

One of the primary mechanisms supporting reef recovery is connectivity of damaged reefs to healthier reefs that can provide a source of larval propagules to promote recovery, therefore understanding connectivity of reef metapopulations and their source and sink dynamics is a key objective for management (18,19). Connectivity is defined here as the exchange of individuals among geographically separated subpopulations that comprise a metapopulation (20). In the context of sessile benthic species, such as corals, population connectivity is predominantly driven by larvae dispersal, though other processes such as spawning strategy and output, predation, available settlement habitat, post-settlement survivorship also influence connectivity (20). Larvae dispersal is a biophysical process and the distance that larvae can disperse is largely dependent on ocean current direction and velocity as well as pelagic larval duration (PLD) and larval competency period (21–23). PLD and larval competency period varies between coral species and can be influenced by environmental conditions such as temperature, which influences developmental rate and time to competency, with increases in SSTs significantly decreasing population connectivity (24,25).

Recovery from major bleaching events has been recorded for remote coral reef systems of the Seychelles, Glorieuses islands and Chagos Archipelago in the western and central Indian Ocean (3,5,8,26,27). In particular, the Chagos Archipelago, highly impacted by the 1998 mass coral bleaching event was reported to have fully recovered its coral cover and structural complexity ten years later (28). These reef systems, which comprise a substantial portion of the western Indian Ocean coral network, are likely linked oceanographically by the South Equatorial Current (SEC) which shows considerable annual and interannual variation in direction and magnitude, but broadly flows westwards from the Indonesian region across the IO to the northern coastline of Madagascar, where it bifurcates at ∼17°S (29,30). The north-flowing branch of the SEC accelerates around Madagascar’s northern tip, generating instabilities that form the Glorioso Front, along with various gyres and mesoscale eddies, which promote high water mixing in the northern Mozambique Channel (31,32).

The south-flowing branch of the SEC flows southward along the east coast of Madagascar before feeding the Agulhas system (33). Seasonally, the SEC and SECC change in intensity for about six months due to varying wind forcing, further influencing circulation patterns across the region (30,33).

The degree of connectivity between coral metapopulations is quantified through population genomic studies or particle tracking simulations based on oceanographic models that provide predictions of larvae dispersal trajectories and population structure. Both methods have their limitations, but integration of these approaches can help explain ocean currents’ influence on connectivity and provide validation for these simulations (34). Examples in the WIO region include recent integration of oceanographic models and population genomic data for the broadcast-spawning coral *Porites cf. lutea*, which suggests high connectivity within the Seychelles (35). Whilst some reef fish species in the Chagos Archipelago have been shown to share population genetic structure with the WIO and the central Indo-Pacific, and there are suggestions that Chagos serves as a genetic ‘stepping-stone’ across the IO (28,36,37)

Corals in the genus *Acropora* dominate on many reefs throughout the Indo-Pacific, and play a key ecological role in reef systems through the provision of habitat structure; however, they are susceptible to a range of disturbances including bleaching from marine heatwaves (13,38). *Acropora tenuis* was considered a widespread and often abundant species on many reefs from the central Pacific to the WIO based on morphological taxonomic work from the late 20^th^ century (38,39). However, recent research incorporating phylogenomic evidence revealed that this putative species was comprised of multiple distinct species with restricted geographical ranges within the Indo-Pacific (40). One of these species, collected from Chagos and Mayotte, did not closely resemble the type material of any nominal species from the region. It was designated as *A.* aff. *tenuis ‘WIO’* in the taxonomic revision of Bridge et al. (40) and is almost certainly undescribed and awaiting formal naming. In this study we investigate the population structure of this species (referred to as *A.* aff. *tenuis* hereafter) across six locations in Chagos, Seychelles and Ile Glorieuses, using target capture of ultraconserved elements (UCEs) and exon loci (41)

UCE and exon loci have the potential to resolve phylogenetic relationships at shallow evolutionary scales because highly divergent and phylogenetically informative regions flank both UCE and exon loci (42) Previous studies have successfully demonstrated that UCE and/or exon data can be used for population genomics (43–45) in a wide range of taxa (e.g., arthropods, frogs, snakes, birds) across a range of evolutionary timescales, including for corals (46). Here we use Cowman *et al.’s* (47) target enrichment of ultra-conserved elements (UCEs) and exons bait set to investigate population structure of *A.* aff*. tenuis* specimens and directionality of gene flow between these populations, as well as to confirm species delimitation. We also run particle tracking simulations to understand the influence of ocean currents on larval dispersal distance and direction and how this correlates with gene flow between *A*. aff. *tenuis* populations, to better understand the drivers of population structure. In addition, we test the impact of positive Indian Ocean Dipole (IOD) phases, which can increase SSTs in the WIO and alter the direction of the SEC (9), on larval dispersal distance and direction and how this correlates with gene flow between *A.* aff*. tenuis* populations.

## Methods

### Field sites

The Chagos Archipelago consists of 2,731km^2^ of reef area across a 60 000km^2^ atoll system (3), and has been a large no-take marine protected area since 2010. There has been no permanent human presence in Chagos since the 1970s except for the US military base on Diego Garcia (48). The Seychelles, more than 1,500 km to the west of Chagos, incorporates 1,935km^2^ of reef area spread across 1.4 million km^2^ of ocean, spanning the Inner granitic islands and Outer coralline islands and including the Aldabra UNESCO world heritage site in the Outer islands (3). The Glorieuses Archipelago, ∼240km SE of Aldabra (Seychelles’ most westerly atoll), is a coral bank 17 km long and covering 165 km² with two main coral islands. Belonging to the Iles Eparses and is administered as part of the TAAF (Terres Australes et Antartiques Françaises), Glorieuses was designated a marine protected area in 2012 and has limited human occupation (49).

For this study, reef sites were grouped into the following six areas based on geographical location; the northern Chagos atolls, western Chagos atolls, Diego Garcia (southern Chagos atoll), Iles Glorieuses, Outer Seychelles Islands and Inner Seychelles Islands. The Chagos Archipelago was grouped into three areas due to its large reef area and our relatively large sampling in this region. Each coral fragment was collected during SCUBA dives at depths of 6 -12m, each colony was photographed and catalogued to verify in-situ species identifications by one of the authors with reference to the material examined in Bridge et al. (40). To avoid collections of genetically identical colonies, fragments were collected from colonies at least three meters distance from each other. Fragments were stored ethanol at -20°C before transportation.

In total, 228 fragments of *A.*aff*.tenuis* were collected from live colonies using pliers to snap off small branch fragments (approx. 1-2cm length) across reef sites in Iles Glorieuses (CITES permit FR1998400003-E), the Outer Seychelles, the Inner Seychelles (CITES permit A1160), Chagos Northern atolls, Chagos Western atolls, and Diego Garcia (Chagos southern atoll) (CITES permits IO/0029/2019, IO/0020/2020, IO/0012/2022) during multiple trips from 2018-2022 (See Figure 1). A total of 217 *A*. aff*. tenuis* specimens were successfully extracted, amplified, and sequenced from across Iles Glorieuses (n=9), Outer Seychelles (n=27), the Inner Seychelles (n=31), Chagos northern atolls (n=105), Chagos western atolls (n=12), and Diego Garcia (n=33) (See Table S1 for reef sites).

**Figure. 1.**
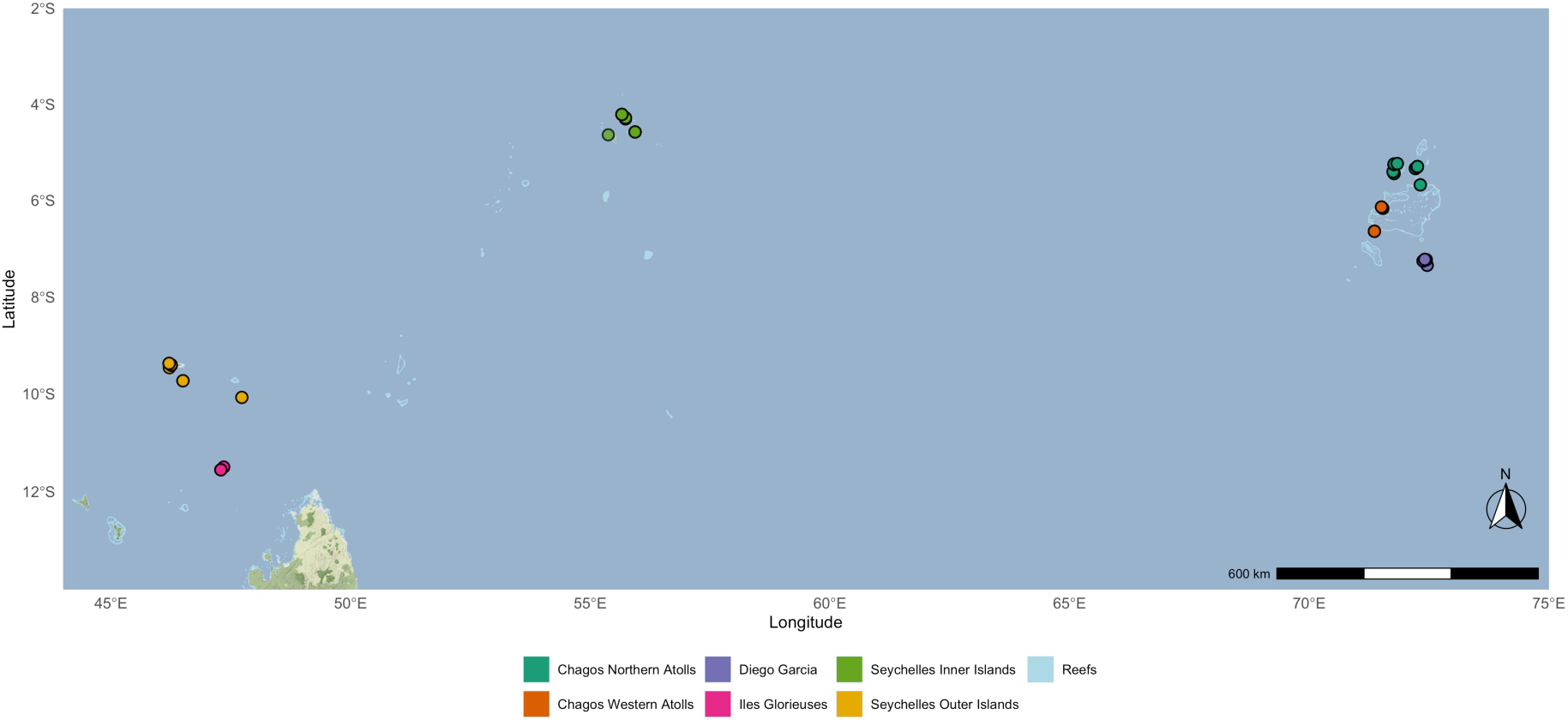
The map gives an overview of all sites which compose the six study locations in the Indian Ocean (see Table S1 for reef sites). Sites are represented by coloured dots and colour-coded by sampling location, and reefs are represented by the light blue colour.

### Extractions, Library-Prep & Sequencing

DNA was extracted with a modified version of the manufacturer protocol for the Qiagen DNeasy PowerSoil kit (Protocols.io: dx.doi.org/10.17504/protocols.io.bww6pfhe). DNA quality was assessed using Qubit fluorometric quantification. Library preparation and target enrichment was performed following methods outlined in Quattrini et al., 2018. An average of 600 ng DNA was sheared to a target size range of 200–400 bp with NEBNext UltraTM II FS DNA Library Prep Kit, successful DNA shearing was assessed with 2% agarose TAE gel. All the sheared DNA was used in downstream library preparation with the Kapa Hyper Prep (Kapa Biosystems). Universal Y-yoke oligonucleotide adapters and custom iTru dual-indexed primers as described by (50) were used for multiplexing. For target enrichment, the *hexa-v2-scleractinia* bait set designed for Scleractinia by Cowman et al., (47) was used. This bait set includes 16,114 baits which are designed to target a total of 2,474 loci (1,127 loci UCE and 1,347 loci exon loci). The Arbor Biosciences MyBaits v. IV protocol was followed, and was performed on pools of up to eight samples. Following target-capture enrichment, target-enriched libraries were sent to Genewiz UK Ltd for sequencing on an Illumina HiSeq 2500 (150 bp PE reads).

### Bioinformatics – Processing, assembly and alignment

The Phyluce pipeline, designed for UCE processing, was used to clean and assemble read data, extract and align loci, and clean these alignments (51). Reads were cleaned using illumiprocessor and trimmomatic v0.35 (52). Contigs were assembled using the program spades v3.12 (53). UCE and exon bait sequences were then separately matched to the assembled contigs at 70% minimum identity and 70% minimum coverage using the command *phyluce_assembly_match_contigs_to_probes*. Loci were then extracted using the command *phyluce_assembly_get_fastas_from_match_counts* and aligned with mafft version 7.130b (54). Edges of the aligned loci were then trimmed using command *phyluce_align_seqcap_align*. Finally, a data matrix including all loci concatenated with at least 75% of taxa present was generated using the command *phyluce_align_get_only_loci_with_min_taxa*.

### SNP Calling

SNP calling was performed using modified scripts from Zarza et al. (45). The coral specimen with the largest number of UCE/exon contigs as identified by command *phyluce_assembly_get_match_counts* was chosen as a reference individual for SNP calling. The commands *phyluce_assembly_get_match_counts* and *phyluce_assembly_get_fastas_from_match_count* were rerun on the reference individual to create a fasta file of UCE and exon contigs found only in the reference. Sequence reads of each sample were mapped to the reference using bwa v 0.7.7 (55) and (56) and removing duplicates using Picard version 1.106-0 (http://broadinstitute.github.io/picard/).

GATK v3.4 (57) was then used to realign BAM files around indels, call variants, and filter variants based on VCF tools v0.1.16 (58) VariantFiltration was implemented for variant quality filtration, and only biallelic SNPs were retained. VCF tools was also used to remove minor allele frequency of 0.05 and genotypes ≤ 10x depth of coverage in each individual. We identified outlier SNPs using bayescan v2.1(59) with 10,000 output iterations and 100 prior odds. The final putative neutral SNPs dataset was built with a missing data rate of less than 10%. Nucleotide diversity (*F*_ST_) for each site was calculated using STACKS (60). All scripts for SNP calling can be found at our Github repository (see data accessibility statement).

### Species verification

Maximum-likelihood (ML) inference was run on the final neutral SNPs dataset to obtain a phylogenetic tree using IQ-Tree2 with 1000 bootstrap searches and 20 ML searches using rapid bootstrapping (61). The best-fitting model of evolution was inferred from the data using IQTREE’s automatic model testing algorithm and was found to be GTR+F+I+R5. The resulting tree was manually mid-rooted, as no outgroup samples were carried through the SNP filtering process and therefore were not available to use to root the tree.

### Population Structure Analysis

To identify clusters of genetically related individuals from the filtered SNPs we used two different approaches. STRUCTURE 2.3.4 (62), which employs Bayesian clustering using a Markov chain Monte Carlo (MCMC) approach, was run under the admixture model to infer the number of genetic clusters (K), testing values from 1 to 7 with prior population information. Analyses included 10 runs with a burn-in length of 100,000 and 1,000,000 MCMC replications. HARVESTER 0.6.94 (63) was used to visualise and analyse the STRUCTURE output by plotting log probability L (*K*) and Δ*K* (64) For each value of *K,* runs were summarized using CLUMPP v.1.1.2 (65). In addition, a discriminant analysis of principal components (DAPC) was also run, using BIC values to find the clusters, in the R package *adegenet* 2.1.10 (66). Finally, to estimate the genetic variance between our sites we calculated pairwise F*st* values using an Analysis of Molecular Variance (AMOVA) approach to partition genetic variance between and within populations, using STACKS v.2.6 (60). All analyses were implemented on the University of Essex CERES server.

### Gene Flow Analysis

Directional relative migration was calculated from allele frequency data using Jost’s D (67) in the R package *diveRsity* (68) using the function divMigrate with 500 bootstraps and a filter threshold of 0.05 (69). The method is based on defining a hypothetical pool of migrants for a given pair of populations and estimating an appropriate measure of genetic differentiation between each of the two populations and the hypothetical pool. The directional genetic differentiation can then be used to estimate the relative levels of migration between the two populations, and the larger of the two relative migration values indicates the population that is likely to behave as a source population (69)

### Oceanographic Modelling

Virtual spawning events were simulated using TrackMPD Lagrangian particle tracking routines (70) that require surface ocean currents, here obtained from the Copernicus Marine Service Global Ocean Physics Reanalysis numerical model. The GLORYS12V1 product (https://doi.org/10.48670/moi-00021) is a global ocean, eddy-resolving NEMO (Nucleus for European Modelling of the Ocean) model with a horizontal resolution of 1/12° on an equirectangular Arakawa C native grid (horizontal resolution ∼9.3 km) and daily temporal resolution.

Particles were released from each of the six locations, and tracked for 120 days, which represents the longest modelled PLD potential for *Acropora* (Connolly & Baird, 2010). Larval competency was assumed to be the same as PLD, however it is acknowledged that larval competency may well be less than the PLD (21,75) but is not known for our study species, and as such our dispersal models represent maximum connectivity. A larval pre-competency window was not incorporated into our model, and again is not known for our study species, but pre-competency has been found to differ between 2-6 days post fertilization within and between species (21). Such small pre-competency windows would likely have little effect on long distance dispersal estimates at distances between locations in this network. Particle releases were spaced at 0.05 degrees in latitude and longitude from pixels containing coral reefs at each location. Therefore, particle releases per location were scaled with geographical expanse of the reef area, defined by the number of pixels covered by the reef. The following number of particles were therefore released per simulation: Iles Glorieuses n= 442, Seychelles Outer Islands n= 886, Seychelles Inner Islands n = 546, Chagos western atolls n = 247, Chagos northern atolls n =299, Diego Garcia n = 117. Corals were assumed to have constant fecundity. Releases were made every two weeks for six months of every year (October – March) between 2010 – 2020 to represent the coral spawning window in the Indian Ocean (76,77) and capture oceanographic conditions in the 8-10 yrs pre - coral collections (equaling approx. 2 generations) when the majority of the corals sampled would have matured, based on colony size at collection, and subsequently begun spawning. Connectivity matrices of the total % of particles passing through the location’s grid cell over the 120day simulation duration were generated at the end of each release period, and a mean connectivity matrix was produced by combining matrices from all years. See supplementary materials for further details and equations.

### Comparisons with oceanographic modelling predictions

A Mantel test was used, in R package vegan (78) to assess correlation between the genetic connectivity matrix between source and sink populations of directional relative migration, and the simulated mean connectivity matrix from the particle tracking models. To test for the effect of positive IOD phases on connectivity, positive IOD years (2015-2016, and 2019-2020) were removed from the particle tracking data set, then a Mantel test was undertaken between this new simulated mean connectivity matrix in negative IOD phases only and the original simulated mean connectivity matrix in both negative and positive IOD phases. In addition, a Mantel test was also undertaken between the genetic connectivity matrix and the simulated mean connectivity matrix in negative IOD phases only.

## Results

A total of 8,220,190 raw reads were generated from the final full dataset of 217 samples. Following quality control and adapter trimming, reads per sample ranged from 1,005,289 to 10,250,647, with a median length of 134 bp. Trimmed reads were assembled into a mean of 84,267 contigs per sample (range: 4,265–701,252) with a mean length of 351bp (range: 78 -12,738bp). The average number of contigs above 1kbp per individual was 2,337 contigs.

The final combined concatenated alignments contained 1,122 UCE/exon loci in the 75% matrix. To discern populations, an initial 14,437 unfiltered SNPs were recovered, and after filtering, 6,424 SNPs remained. Based on testing with BAYESCAN, no SNPs were found to be under selection. The tree topology, resulting from Maximum-likelihood inference, provided reassurance on the phylogenetic distribution of the sample set (Fig. S1).

### Genetic Population Structure

STRUCTURE clustering revealed that the mean likelihood probability for the values of *K,* ranging from 2 to 7, plateaued at K=2 (Fig.S2), therefore the most likely number of genetic clusters is two (Fig. 2). Clustering patterns for all K values can be seen in Fig. S3. One of these genetic clusters was found across all locations, dominating the Chagos Archipelago and Inner Seychelles with few exceptions, and occurring only occasionally in Iles Glorieuses, where evidence of introgression from this genetic cluster was minimal. Iles Glorieuses was the only site where individuals primarily belonged to the rarer second cluster (Figs. 2). In contrast, the Outer Seychelles showed mixed ancestry from both clusters. The rarer cluster was detected infrequently in the northern Chagos atolls and was nearly absent from the western and southern Chagos and the Inner Seychelles (Fig. 2). The Discriminant Analysis of Principal Components (DAPC) also shows the same pattern, with two clusters most strongly statistically supported by BIC (Fig.S4). The DAPC visualization, with pre- defined populations, also reveals finer-scale structure consistent with 3 broader subgroupings (Fig. S5). DA1 and DA2 explain 58.4% and 22.0% of the between-group variance, respectively. Finally, to estimate the genetic variance between our sites we calculated pairwise F*st* values; low F*st* values (> 0.1) were estimated for all combinations (Table S2).

**Figure 2.**
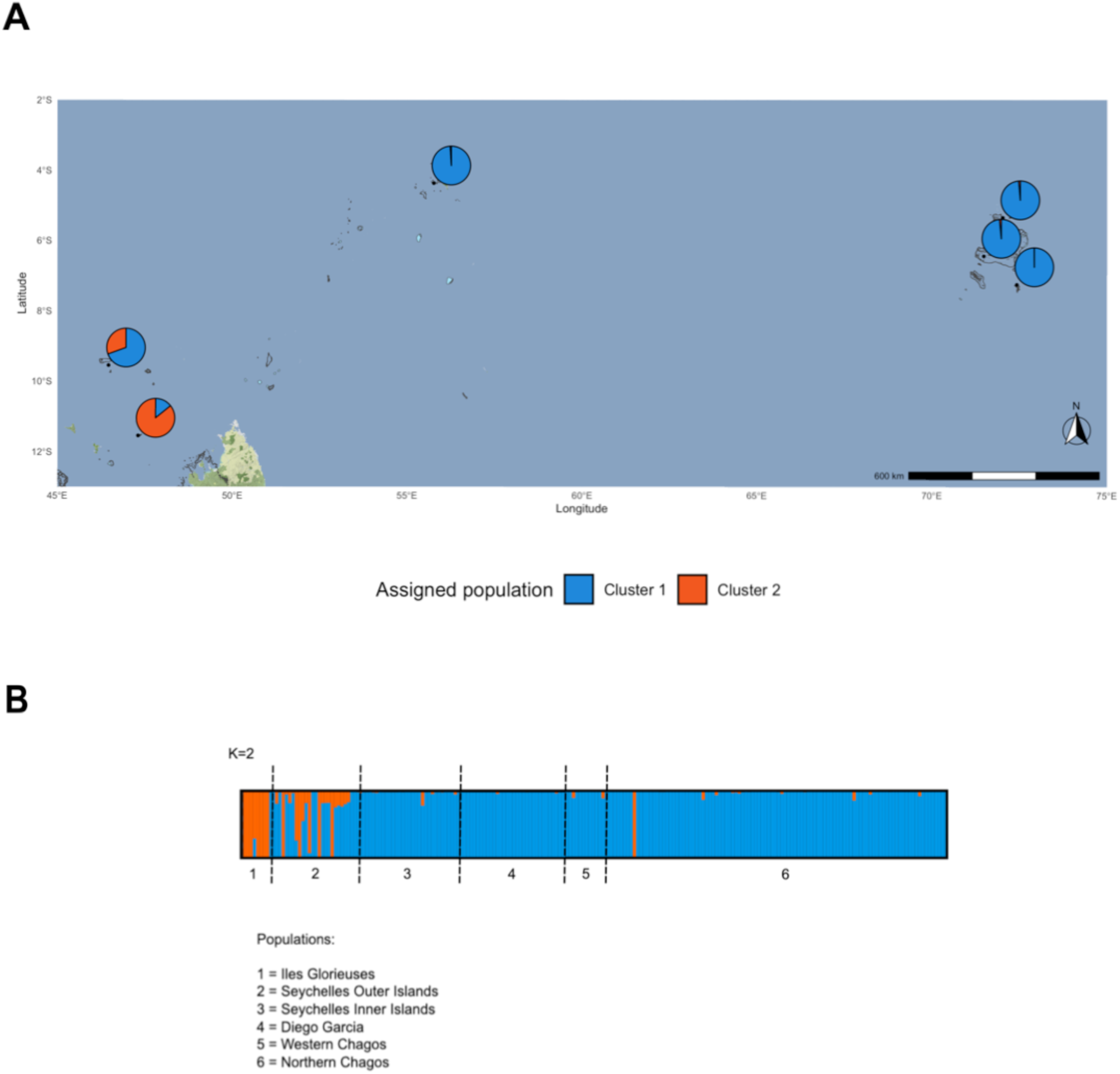
Population structure of *A.* aff. *tenuis* for K=2 for the six study locations, showing two lineages. (A) represents the population structure of each location as a pie chart on the map of the study area (background lines represent reefs). (B) represents the population structure in a barplot.

### Gene Flow

Relative migration/gene flow occurs between all locations but is highly asymmetrical (Fig. 3). Migration from Iles Glorieuses to other sites was significantly higher than in the opposite direction (Fig. 3 & Table S3), suggesting these sites act as source reefs. However, overall migration rates from Iles Glorieuses were an order of magnitude lower than between most other locations, indicating limited connectivity. Conversely, migration into the Outer Seychelles from all other sites was significantly higher than the reverse, suggesting these reefs function as sink reefs. Migration rates from Diego Garcia to Northern Chagos, and from Inner Seychelles to Diego Garcia are also significantly higher relative to the opposite direction, but gene flow is high in both directions in both cases (Fig. 3 & Table S3). Reciprocal gene flow among the three Chagos locations is strong, as is connectivity between the Inner Seychelles and the Chagos sites, indicating high regional connectivity (Fig. 3).

**Figure 3.**
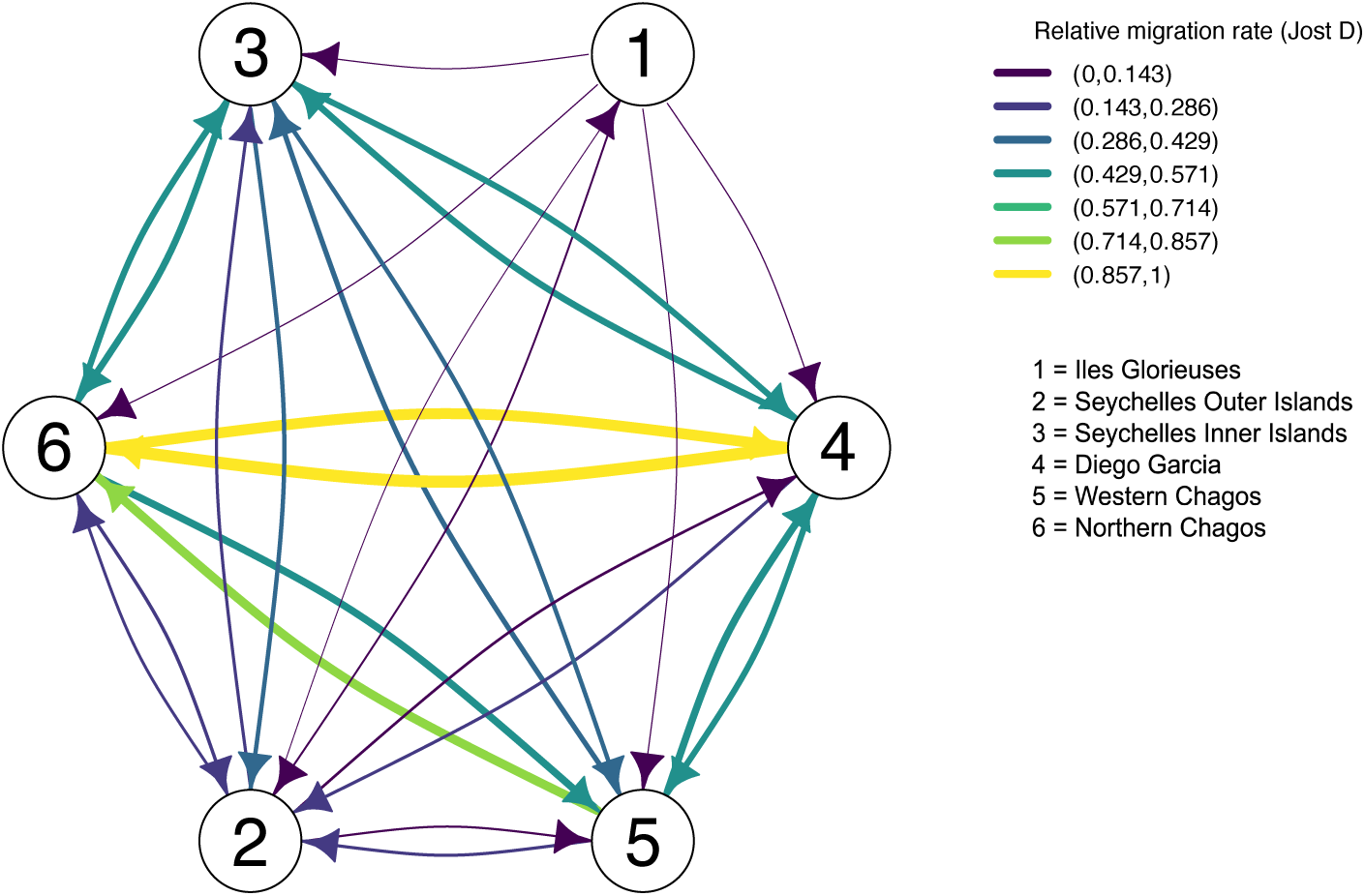
Relative migration network of the six study locations demonstrate patterns of connectivity and geneflow. Directionality of arrows represents direction of gene flow. Each arrow is colour coded to show the strength of gene flow measured by Jost’s D relative migration rate metric. The relative migration rate (Jost’s D) statistics can also be found in Table S3.

### Comparisons with oceanographic modelling predictions

Lagrangian particle tracking models were summarised into a connectivity matrix, calculated as the percentage of particles from one location passing through the area defined as another of the six locations across the whole 120 day duration of the simulations (Fig. 4). Mantel tests showed no significant correlation between distance connectivity matrices, measured by gene flow (Jost’D metric of genetic differentiation), and measured by particle tracking models (number of particles passing through that location’s grid cell(s)) (*r = −0.164 p= 0.654*). Therefore, as the distance between locations increases with respect to the genetic connectivity matrix, the distances between the same locations do not necessarily increase in the particle tracking connectivity matrix.

**Figure 4.**
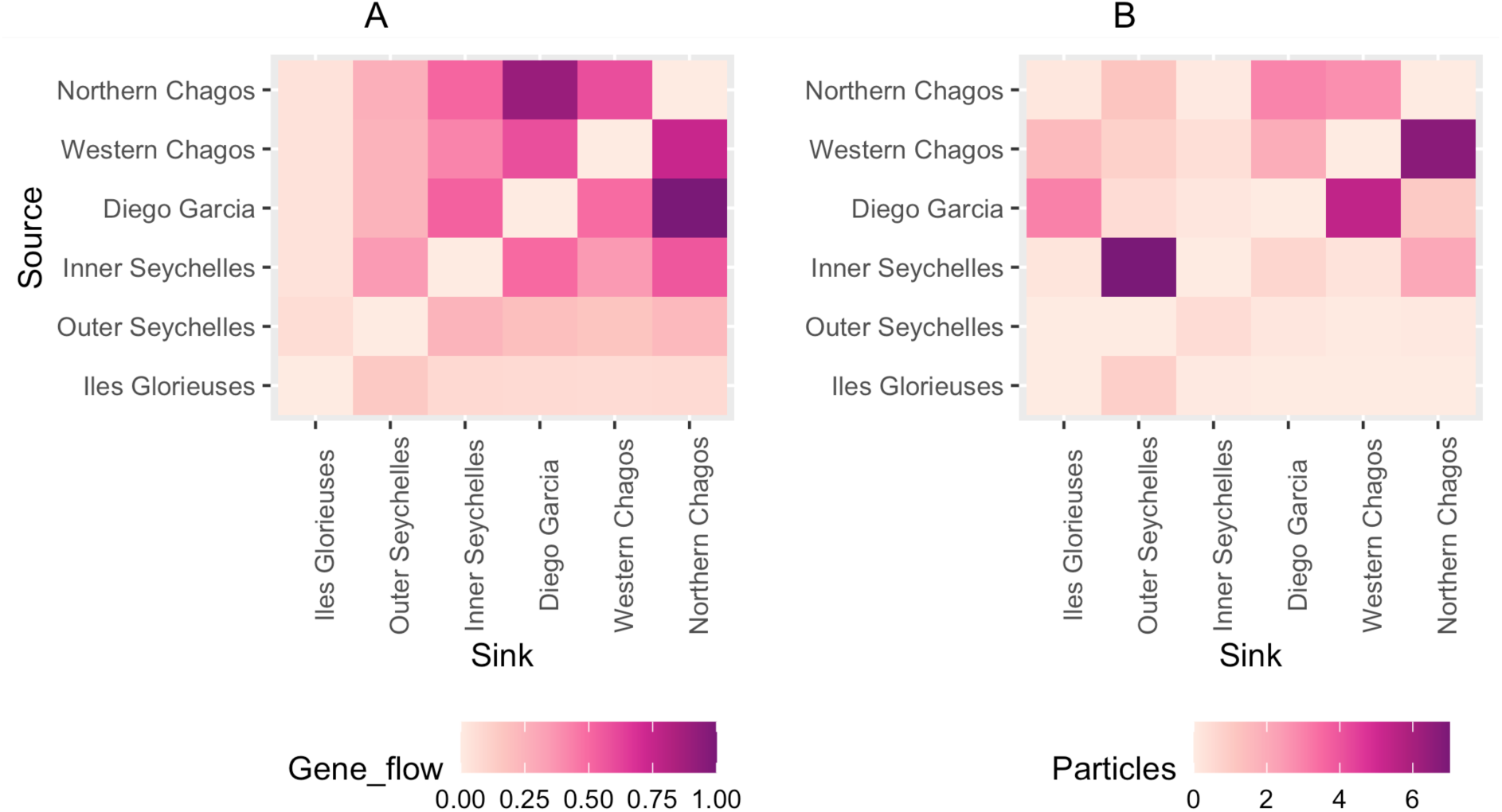
Heat maps representing A) the genetic connectivity measured by gene flow (migration rates) between source and sink locations, B) the simulated connectivity measured by the % of particles passing through the grid cell from source to sink locations. Darker colours represent stronger connectivity.

### Comparisons with oceanographic modelling predictions in negative IOD phases only

Mantel tests showed that there was a significant correlation between distance connectivity matrices measured by the particle tracking models in years with both positive and negative IOD phases and models with just negative IODs phases (r= 0.607, p= 0.015), suggesting that the pattern of dispersal is not significantly changed by the change in IODs. However, the connectivity means are generally higher in models without positive IODs, suggesting that positive IODs weaken connectivity between these locations (Fig S6 illustrates particle movement in +ve and -ve IOD years)). However, Mantel tests examining the correlation between connectivity from particle tracking models in negative IOD years only and gene flow connectivity, showed no significant change in the correlation patterns from tests of models with both negative and positive IOD years (r = −0.214, p=0.715, and *r = −0.164 p= 0.654* respectively).

## Discussion

In this study we identify widespread connectivity in *Acropora* aff. *tenuis* across remote atolls in the central-western Indian Ocean, a finding with positive implications for the capacity of these reefs to recovery from disturbance such as bleaching events. Gene flow was particularly strong within and between Chagos and the Inner Seychelles, indicating a remarkable larval supply network operating between these atolls, especially considering the straight-line distances of over 1500 km. This is congruent with strong asymmetrical larval dispersal predictions between Chagos and the Inner Seychelles (33). Larvae transport between the Inner and Outer Seychelles has recently also been found to be substantial for other coral taxa (35). In our study, gene flow between Chagos and Inner Seychelles appears stronger than between the Outer and Inner Seychelles, which are geographically closer. This may be partially explained by a possible permeable barrier to dispersal created by ocean currents between the Outer and Inner Seychelles island groups (35). This is further supported by the population structure of the rarer of the two *A.* aff. *tenuis* linages in our study, which appear more frequently in Iles Glorieuses and in the Outer Seychelles reefs, the two more westerly locations, but are rare in the more easterly locations.

Directionality in migration rates suggest the Outer Seychelles is a sink reef in this network, receiving significantly more larvae from Chagos and the Inner Seychelles than it provides. This is likely as a result of the prevailing current, which largely results in larvae from the outer islands traveling westward towards the East African coastline (33) In contrast, Iles Glorieuses is acting as a source reef in this network; however, migrations rates were relatively low, suggesting it is the most isolated of all the atolls in the network despite its relatively close proximity to the Outer Seychelles. Iles Glorieuses’ position in the Mozambique channel mean it is subjected to complex oceanographic patterns of eddies and gyres. These oceanographic features increase random variation in larvae dispersal patterns, which likely have greater influence on connectivity than geographical distance (79) and may explain these low migration rates. Reef area is also a factor that limits potential larval supply from source reefs, less coral means less larvae to disperse (80), and comparatively, Iles Glorieuses’ reef area is significantly smaller than that of the Seychelles and Chagos.

The population structure of *A.*aff *tenuis* comprised two distinct lineages, with geographical mixing between the two lineages most evident in the Outer Seychelles. The spatial distribution of these two linages may be a result of a founder effect, which occurs when only a small number of larvae from one or two spawning events colonizes a new reef, leading to a loss of genetic variation and the establishment of a founder population with a distinct genetic composition ((81,82). Small, isolated populations, such as those on many remote atolls, are more likely to exhibit genetic drift, leading to this phenomenon. There is evidence that a small proportion of coral spawning events account for the majority of settling larvae in the Indian Ocean system supporting this idea (83). When a founder population is established from a small number of individuals, it is likely to exhibit reduced genetic diversity and as a result, founder populations may be less able to adapt to environmental changes (82).

The Outer Seychelles, as a sink population with evidence of both lineages, may represent a population with a higher genetic diversity, and as such a greater evolutionary advantage to adapting resilience mechanisms to environmental changes, such as marine heatwaves.

On the coast of mainland Africa and Madagascar, to the west and south west of our study sites, Van der ven et al. (84,85) identified five lineages of *Acropora* cf. *tenuis* (identified as *A. tenuis* but now known to be outside the geographic range of *A. tenuis* which occurs only in the south-west Pacific). Irrespective of the taxonomic identity, the species examined showed high but variable genetic connectivity between East African populations, driven by the prevailing northbound East African Coast Current, and high differentiation between the African mainland and Madagascar and within Madagascar. Despite high genetic differentiation there was evidence for long-distance larval dispersal for *A.* cf. *tenuis* across the North Mozambique Channel region, with exchange between northern Mozambique and northern Madagascar. The occurrence of only two linages in our study may suggest limited long-distance dispersal from Iles Glorieuses to the west, or that Van der ven et al. examined multiple morphologically similar species. However, it is difficult to make inferences as our studies used different genetic approaches and the *A. tenuis complex* in the WIO is yet to be fully resolved.

### Drivers of Connectivity

Ocean currents are a major driver of coral larvae dispersal mainly because of coral larvae’s poor swimming ability (86) The strong connectivity between Chagos and the Inner Seychelles is likely due to the eastward South Equatorial Countercurrent (SECC) influencing the Inner Islands of Seychelles during the northwest monsoon, providing an efficient pathway for eastward larval dispersal towards the Chagos Archipelago, and in the other direction the predominant westward flowing SEC also during the spawning season (October-April) facilitates larvae transport from Chagos to the Inner Seychelles Islands and to all other locations (33). These powerful current systems between Chagos and the Inner Islands of the Seychelles suggest that regular connectivity is possible and may shape the strong genetic connectivity between these locations, rather than chance events in certain years driven by idiosyncratic variations in oceanographic conditions.

High gene flow within Chagos is likely a result of its expansive area of reef cover resulting in a relatively high quantity of larvae production, as well as relatively weak currents within much of the Archipelago encouraging larvae retention (33,80) Larvae retention, and subsequent self-recruitment of reefs, is known to be critical for the recovery and resilience of isolated reef systems (87,88), and maybe largely responsible, alongside the absence of many direct anthropogenic stressors, for Chagos’ strong recovery from the 1998 bleaching event (28). Given Chagos’ ability to act as a source reef to the WIO through the predominant westward flowing SEC during the spawning season (33), protecting these reefs and those of the Inner Seychelles, with which Chagos is strongly connected, may be of high value to the persistence and recovery of the wider IO reefs from global climate change.

The lack of significant correlation between connectivity derived from the particle tracking models and from genetic migration rates indicates that large-scale ocean currents are not the only driver of larvae dispersal across the region. The temporal resolution of oceanographic models does not always accurately capture the stochastic short-term (e.g. daily) variability in oceanographic processes, which can limit the accuracy of connectivity models (83,89). Submesoscale oceanographic processes and reef-scale hydrodynamics (1 -1,000m) (Fig. S7), which cannot be captured at the resolution of global scale oceanographic models (100m – 10,000m), are powerful drivers of genetic connectivity (90,91). Given that small-scale processes influence local retention rates and whether larvae in the open ocean arrive at nearshore reef systems, quantifying these processes is important for understanding regional connectivity patterns.

These difficulties in capturing small scale processes are also exacerbated by uncertainty in larval development, behaviour, and mortality, as well as adult spawning behavior (20,21,75,80) Long- distance dispersal is strongly influenced by reproductive output (abundance and fecundity of adults) and length of the PLD, whereas self-recruitment is primarily driven by local oceanography, larval mortality and the larval pre-competency window (80) Meanwhile, duration of spawning has recently been identified as a major driver of connectivity rates (83). Relatively little is known about coral spawning in most species (76) with published knowledge on spatial and temporal spawning patterns from just a few species and locations in the IO (77,92–94). Therefore, while the timings and duration of particle releases in our models are based on the best available information they may not be entirely accurate for the study species, and larval dispersal predictions are known to be highly sensitive to these hydrodynamic model choices (21,89,95). Preliminary data on the *A.cytherea* complex *(in prep.)* suggests much more limited connectivity across the same network than for *A .*aff. *tenuis*, demonstrating that coral reproduction biology, even within broadcast spawners, can differ substantially between species and may be more influential on connectivity than previously considered (21,96)

### Indian Ocean Dipole (IOD) phases

We also find that positive IOD events (2015-2016, and 2019-2020) caused a weakening in simulated connectivity, but not significantly so. This weakening may be caused by increased meandering of the SEC in positive IOD years (30) and the associated rise in upper ocean temperature in the WIO leading to an increased larval mortality rate and more rapid larval development rates reducing connectivity (97,98). As global climate change impacts continue to increase we may see larger variation in IOD patterns in the WIO (99), with a knock-on effect on connectivity and larval supply.

### Stepping-stone-effect

The strong genetic connectivity across these remote reef systems is also likely a result of a stepping-stone-effect (expected dispersal over multiple generations) across locations. The large geographic distance between the westernmost atoll (Iles Glorieuses) and the easternmost (Chagos) is >2,500 km, a distance that is likely to rarely, if ever, be covered by coral larvae. Stepping stones are often crucial to long-distance dispersal (100) therefore incorporating a comparison of multi-generational dispersal from biophysical models can improve the fit with spatial genetic structure (34). Given that *Acropora* generally mature within 5 years and our simulations spanned 10 years (2 generations), it was not possible to examine the importance of stepping stones across multiple generations. However, recent multidecadal larval dispersal models considering multi-generational dispersal indicate that Chagos, the Seychelles and three small island groups in the path of the SEC (the Agalega Islands, St Brandon, and Tromelin) physically share ancestral populations within 10-100 dispersal steps, equivalent to around 50 - 500 years (33). Importantly, this suggests that regional population structure can be attributed to evolutionary recent dispersal events (33).

### Conclusion

This study advances understanding of reef connectivity among geographically remote atolls in the Indian Ocean, providing evidence of long-distance larval dispersal across a reef network spanning vast distances. Strong connectivity between Chagos and the Inner Seychelles may be especially important for regional management. Continued protection of Chagos reefs is likely to support reef resilience in the Inner Seychelles via larval supply, where local communities depend heavily on reefs for tourism and fisheries, as well as in the wider region via this stepping-stone connectivity pathway. In contrast, the Outer Seychelles appear to function as a sink population, hosting both *A.* aff*. tenuis* lineages, which may enhance resilience to future environmental stressors such as marine heatwaves. Yet, identifying which reef locations are most likely to survive under continued warming across complex seascapes like the western Indian Ocean remains a major challenge. Understanding how connectivity interacts with thermal tolerance is a key component of adaptive management in the face of rapid environmental change (101). Remote atolls with minimal direct human impact, such as Chagos, the Outer Seychelles, and Iles Glorieuses, may play a critical role in maintaining regional reef resilience through their contributions to long-distance dispersal.

## Ethics

CITES permits were obtained for coral specimen collection from each location; Iles Glorieuses (CITES permit FR1998400003-E), the Outer Seychelles and the Inner Seychelles (CITES permit A1160), Chagos Northern atolls, Chagos Western atolls, and Diego Garcia (Chagos southern atoll) (CITES permits IO/0029/2019, IO/0020/2020, IO/0012/2022) during multiple trips from 2018-2022.

## Declaration of AI Use

AI-assisted technologies were used on occasion to aid in identification of suitable code for analysis.

## Acknowledgements

This work was supported by the Bertarelli Foundation as part of the Bertarelli Programme in Marine Science (Chagos Science Expeditions 2018, 2019, 2020). We thank the captain and crew of the Chagos patrol vessel for their logistical support. We also thank the Monaco Explorations and Prince Albert II of Monaco Foundation for berths on the Scattered Island Expedition 2019. We thank the many in-kind contributions we received to facilitate transport and diving during fieldwork in the Seychelles including site access and permissions from the Seychelles Islands Foundation, Fregate Island Conservation, Le Ponant, Silverseas, Seychelles National Parks Authority, Island Conservation Society Seychelles and the Marine Conservation Society Seychelles. Funding for the Seychelles part of this study was supported by the NERC environmental research doctoral training programme at the University of Oxford (Grants NE/L002612/1) with additional funding by the Seychelles Climate Change Adaptation Trust (SeyCCAT). We thank Research England for supporting CY and CH.

## Bibliography

1. Hughes TP, Kerry JT, Álvarez-Noriega M, Álvarez-Romero JG, Anderson KD, Baird AH, et al. Global warming and recurrent mass bleaching of corals. Nature. 2017 Mar 15;543(7645):373–7. doi:10.1038/nature21707 PubMed PMID: 28300113.

2. Hoegh-Guldberg O, Skirving W, Dove SG, Spady BL, Norrie A, Geiger EF, et al. Coral reefs in peril in a record-breaking year. Science (1979). 2023 Dec 1;382(6676):1238–41. doi:10.1126/science.adk4532 PubMed PMID: 38060674.

3. Souter D, Planes S, Wicquart J, Obura D, Staub F. Status of Coral Reefs of the World: 2020. 2021 Oct. doi:10.59387/WOTJ9184

4. Eakin CM, Heron SF, Connolly SR, Devotta DA, Liu G, Geiger EF, et al. Severe and widespread coral reef damage during the 2014-2017 Global Coral Bleaching Event. Nat Commun. 2026 Feb 10;17(1):1318. doi:10.1038/s41467-025-67506-w

5. Head CEI, Bayley DTI, Rowlands G, Roche RC, Tickler DM, Rogers AD, et al. Coral bleaching impacts from back-to-back 2015-2016 thermal anomalies in the remote central Indian Ocean. Coral Reefs. 2019 Aug 15;38(4):605–18. doi:10.1007/s00338-019-01821-9

6. Cerutti JMB, Burt AJ, Haupt P, Bunbury N, Mumby PJ, Schaepman-Strub G. Impacts of the 2014–2017 global bleaching event on a protected remote atoll in the Western Indian Ocean. Coral Reefs. 2020 Feb 1;39(1):15–26. doi:10.1007/s00338-019-01853-1

7. Koester A, Migani V, Bunbury N, Ford A, Sanchez C, Wild C. Early trajectories of benthic coral reef communities following the 2015/16 coral bleaching event at remote Aldabra Atoll, Seychelles. Sci Rep. 2020 Dec 1;10(1). doi:10.1038/s41598-020-74077-x PubMed PMID: 33046828.

8. Sheppard C, Sheppard A, Fenner D. Coral mass mortalities in the Chagos Archipelago over 40 years: Regional species and assemblage extinctions and indications of positive feedbacks. Mar Pollut Bull. 2020 May 1;154. doi:10.1016/j.marpolbul.2020.111075 PubMed PMID: 32319906.

9. Shi W, Wang M. A biological Indian Ocean Dipole event in 2019. Sci Rep. 2021;28(11):2452.

10. Van Hooidonk R, Maynard J, Tamelander J, Gove J, Ahmadia G, Raymundo L, et al. Local-scale projections of coral reef futures and implications of the Paris Agreement. Sci Rep. 2016 Dec 21;6. doi:10.1038/srep39666 PubMed PMID: 28000782.

11. Van Hooidonk R, Maynard J, Grimsditch G, Williams G, Tamelander J, Gove J, et al. UNEP 2020. Projections of future coral bleaching conditions using IPCC CMIP6 models: climate policy implications, management applications, and Regional Seas summaries. United Nations Environment Programme, Nairobi, Kenya. 2020. Available from: http://www.unep.org/publications/

12. Lachs L, Bozec YM, Bythell JC, Donner SD, East HK, Edwards AJ, et al. Natural selection could determine whether *Acropora* corals persist under expected climate change. Science. 2024 Nov 28. doi:10.1126/science.adl6480

13. Hughes TP, Kerry JT, Baird AH, Connolly SR, Dietzel A, Eakin CM, et al. Global warming transforms coral reef assemblages. Nature. 2018 Apr 26;556(7702):492–6. doi:10.1038/s41586-018-0041-2 PubMed PMID: 29670282.

14. Pearce-Kelly P, Altieri AH, Bruno JF, Cornwall CE, McField M, Muñiz-Castillo AI, et al. Considerations for determining warm-water coral reef tipping points. Earth Syst Dynam. 2025;16:275–92.

15. Boström-Einarsson L, Babcock RC, Bayraktarov E, Ceccarelli D, Cook N, Ferse SCA, et al. Coral restoration – A systematic review of current methods, successes, failures and future directions. PLoS One. 2020 Jan 30;15(1):e0226631. doi:10.1371/journal.pone.0226631

16. Mcleod E, Anthony KRN, Mumby PJ, Maynard J, Beeden R, Graham NAJ, et al. The future of resilience-based management in coral reef ecosystems. J Environ Manage. 2019 Mar;233:291–301. doi:10.1016/j.jenvman.2018.11.034

17. Anthony KRN, Marshall PA, Abdulla A, Beeden R, Bergh C, Black R, et al. Operationalizing resilience for adaptive coral reef management under global environmental change. Glob Chang Biol. 2015 Jan 5;21(1):48–61. doi:10.1111/gcb.12700

18. McManus LC, Forrest DL, Tekwa EW, Schindler DE, Colton MA, Webster MM, et al. Evolution and connectivity influence the persistence and recovery of coral reefs under climate change in the Caribbean, Southwest Pacific, and Coral Triangle. Glob Chang Biol. 2021 Sep 1;27(18):4307–21. doi:10.1111/gcb.15725 PubMed PMID: 34106494.

19. Palumbi SR. Population genetics, demographic connectivity, and the design of marine reserves. Ecol Appl. 2003;13:146–58.

20. Cowen RK, Sponaugle S. Larval dispersal and marine population connectivity. Annual Review of Marine Science. 2009. p. 443–66. doi:10.1146/annurev.marine.010908.163757 PubMed PMID: 21141044.

21. Randall CJ, Giuliano C, Stephenson B, Whitman TN, Page CA, Treml EA, et al. Larval precompetency and settlement behaviour in 25 Indo-Pacific coral species. Commun Biol. 2024 Dec 1;7(1). doi:10.1038/s42003-024-05824-3 PubMed PMID: 38297134.

22. Monismith SG, Barkdull MK, Nunome Y, Mitarai S. Transport Between Palau and the Eastern Coral Triangle: Larval Connectivity or Near Misses. Geophys Res Lett. 2018 May 28;45(10):4974–81. doi:10.1029/2018GL077493

23. Shanks A, Grantham B, Carr M. Propagule dispersal and the size and spacing of marine reserves. Ecol Appl. 2003;13:159–69.

24. Figueiredo J, Baird AH, Connolly SR. Synthesizing Larval Competence Dynamics and Reef- Scale Retention Reveals a High Potential for Self-recruitment in Corals. Ecology. 2013;94:650–9.

25. O’Connor MI, Bruno JF, Gaines SD, Halpern BS, Lester SE, Kinlan BP, et al. Temperature control of larval dispersal and the implications for marine ecology, evolution, and conservation. Proceedings of the National Academy of Sciences. 2007 Jan 23;104(4):1266–71. doi:10.1073/pnas.0603422104

26. Graham NAJ, Jennings S, MacNeil MA, Mouillot D, Wilson SK. Predicting climate-driven regime shifts versus rebound potential in coral reefs. Nature. 2015 Feb 5;518(7537):94–7. doi:10.1038/nature14140 PubMed PMID: 25607371.

27. Gudka M, Obura D, Mbugua J, Ahamada S, Kloiber U, Holter T. Participatory reporting of the 2016 bleaching event in the Western Indian Ocean. Coral Reefs. 2020 Feb 1;39(1):1–11. doi:10.1007/s00338-019-01851-3

28. Sheppard C, Ateweberhan M, Bowen BW, Carr P, Chen CA, Clubbe C, et al. Reefs and islands of the Chagos Archipelago, Indian Ocean: why it is the world’s largest no-take marine protected area. Aquatic Conserv: Mar Freshw Ecosyst. 2012;232–61.

29. Obura D. The Diversity and Biogeography of Western Indian Ocean Reef-Building Corals. PLoS One. 2012 Sep 19;7(9). doi:10.1371/journal.pone.0045013 PubMed PMID: 23028737.

30. Chen G, Han W, Wang D, Zhang L, Chu X, He Y, et al. Seasonal structure and interannual variation of the South Equatorial Current in the Indian Ocean. J Geophys Res Oceans. 2022;127(11).

31. Belkin I, Cornillon P, Sherman K. Fronts in Large Marine Ecosystems. Progress in Oceanography. 2009;81(1–4):223–36.

32. Schouten W, de Ruijter W, van Leeuwen P, Ridderinkhof H. Eddies and variability in the Mozambique Channel. Deep Sea Research, Part II. 2003;50:1987–2003.

33. Vogt-Vincent NS, Burt AJ, van der Ven RM, Johnson HL. Coral reef potential connectivity in the southwest Indian Ocean. Coral Reefs. 2024. doi:10.1007/s00338-024-02521-9

34. Jahnke M, Jonsson PR. Biophysical models of dispersal contribute to seascape genetic analyses. Philosophical Transactions of the Royal Society B: Biological Sciences. Royal Society Publishing; 2022. doi:10.1098/rstb.2021.0024 PubMed PMID: 35067094.

35. Burt A, Vogt-Vincent N, Johnson H, Sendell-Price A, Kelly S, Clegg SM, et al. Integration of population genetics with oceanographic models reveals strong connectivity among coral reefs across Seychelles. Sci. Rep. 2024;14:4936. 10.1038/s41598-024-55459-x

36. Bowen B, Bass A, Rocha L, Grant W, Robertson D. Phylogeography of the trumpetfishes (Aulostomus): ring species complex on a global scale. Evolution. 2001;55:1029–39.

37. Gaither M, Bowen B, Bordenave T, Rocha L, Newman S, Gomez J, et al. Phylogrography of the reef fish Cephalophis argus (Epinephelidae) indicates Pleistocene isolation across the Indo-Pacific Barrier with contemporary ovelap in the Coral Triangle. BMC Evol Biol. 2011;11:189.

38. Wallace C. Staghorn corals of the world: a revision of the genus Acropora. CSIRO Publishing, Collingwood, VIC, Australia.; 1999.

39. Veron JEN. Corals of the World. Vols 1–3. Australian Institute of Marine Science, Townville, 1382; 2000.

40. Bridge TCL, Cowman PF, Quattrini AM, Bonito VE, Sinniger F, Harii S, et al. A tenuis relationship: traditional taxonomy obscures systematics and biogeography of the ‘ Acropora tenuis ’ (Scleractinia: Acroporidae) species complex. Zool J Linn Soc. 2024;202.1–24. doi:10.1093/zoolinnean/zlad062

41. Quattrini A, Faircloth B, Dueñas L, et al. Universal target-enrichment baits for anthozoan (Cnidaria) phylogenomics: new approaches to long-standing problems. Mol Ecol Resour. 2018;18:281–95.

42. Faircloth BC, McCormack JE, Crawford NG, Harvey MG, Brumfield RT, Glenn TC. Ultraconserved elements anchor thousands of genetic markers spanning multiple evolutionary timescales. Syst Biol. 2012 Oct;61(5):717–26. doi:10.1093/sysbio/sys004 PubMed PMID: 22232343.

43. Oswald JA, Harvey MG, Remsen RC, Foxworth DU, Dittmann DL, Cardiff SW, et al. Evolutionary dynamics of hybridization and introgression following the recent colonization of Glossy Ibis (Aves: Plegadis falcinellus) into the New World. Mol Ecol. 2019;28(7):1675–91.

44. Winker K, Glenn TC, Faircloth BC. Ultraconserved elements (UCEs) illuminate the population genomics of a recent, high-latitude avian speciation event. PeerJ. 2018;2018(10). doi:10.7717/peerj.5735

45. Zarza E, Faircloth BC, Tsai WL, Bryson RW, Klicka J, McCormack JE. Hidden histories of gene flow in highland birds revealed with genomic markers. Mol Ecol. 2016;25(20):5144–57.

46. Erickson KL, Pentico A, Quattrini AM, McFadden CS. New approaches to species delimitation and population structure of anthozoans: Two case studies of octocorals using ultraconserved elements and exons. Mol Ecol Resour. 2021 Jan 1;21(1):78–92. doi:10.1111/1755-0998.13241 PubMed PMID: 32786110.

47. Cowman PF, Quattrini AM, Bridge TCL, Watkins-Colwell GJ, Fadli N, Grinblat M, et al. An enhanced target-enrichment bait set for Hexacorallia provides phylogenomic resolution of the staghorn corals (Acroporidae) and close relatives. Mol Phylogenet Evol. 2020 Dec 1;153. doi:10.1016/j.ympev.2020.106944 PubMed PMID: 32860973.

48. Hays GC, Koldewey HJ, Andrzejaczek S, Attrill MJ, Barley S, Bayley DTI, et al. A review of a decade of lessons from one of the world’s largest MPAs: conservation gains and key challenges. Marine Biology. 2020. doi:10.1007/s00227-020-03776-w

49. Quétel C, Marinesque S, Ringler D, Fillinger L, Changeux T, Marteau C, et al. Iles Eparses (SW Indian Ocean) as reference ecosystems for environmental research. Acta Oecologica. 2016 Apr 1;72:1–8. doi:10.1016/j.actao.2015.12.012

50. Glenn TC, Nilsen R, Kieran TJ, Finger JW, Pierson TW, Bentley KE, et al. Adapterama I: Universal stubs and primers for thousands of dual-indexed Illumina libraries (iTru & iNext). PeerJ. 2016. PMCID: PMC6791352 DOI: 10.7717/peerj.7755

51. Faircloth BC. PHYLUCE is a software package for the analysis of conserved genomic loci. Bioinformatics. 2016;32:786–8.

52. Bolger AM, Lohse M, Usadel B. Trimmomatic: A flexible trimmer for Illumina sequence data. Bioinformatics. 2014 Aug 1;30(15):2114–20. doi:10.1093/bioinformatics/btu170 PubMed PMID: 24695404.

53. Bankevich A, Nurk S, Antipov D, Gurevich AA, Dvorkin M, Kulikov AS, et al. SPAdes: a new genome assembly algorithm and its applications to single-cell sequencing. J Comput Biol. 2012;19(5):455–77. doi: 10.1089/cmb.2012.0021.

54. Katoh K, Misawa K, Kuma KI, Miyata T. MAFFT: a novel method for rapid multiple sequence alignment based on fast Fourier transform. Nucleic Acids Res. 2002 Jul 15;30(14):3059–66. doi: 10.1093/nar/gkf436.

55. Li H, Durbin R. Fast and accurate long-read alignment with Burrows–Wheeler transform. Bioinformatics. 2010;26(5):589–96.

56. Li H, Handsaker B, Wysoker A, Fennell T, Ruan J, Homer N, et al. The Sequence Alignment/Map format and SAMtools. Bioinformatics. 2009 Aug 15;25(16):2078–9. doi:10.1093/bioinformatics/btp352 PubMed PMID: 19505943.

57. Van der Auwera G, O’Connor B. Genomics in the Cloud: Using Docker, GATK, and WDL in Terra. 1st Edition. O’Reilly Media. 2020.

58. Danecek P, Auton A, Abecasis G, Albers C, Banks E, DePristo M, et al. The variant call format and VCFtools. Bioinformatics. 2011;27(15):2156–8.

59. Foll M, Gaggiotti O. A genome-scan method to identify selected loci appropriate for both dominant and codominant markers: a Bayesian perspective. Genetics. 2008;180(2):977–83.

60. Catchen J, Hohenlohe P, Bassham S, Amores A, Cresko W. Stacks: an analysis tool set for population genomics. Mol Ecol. 2013;22(11):3124–40. doi: 10.1111/mec.12354.

61. Minh B, Schmidt H, Chernomor O, Schrempf D, Woodhams M, von Haeseler A, et al. IQ- TREE 2: New models and efficient methods for phylogenetic inference in the genomic era. Mol Biol Evol. 2020;37:1530–4.

62. Pritchard M, Stephens J, Donnelly P. Inference of population structure using multilocus genotype data. Genetics. 2000;155:945–59.

63. Earl D, von Holdt B. STRUCTURE HARVESTER: A Website and Program for Visualizing STRUCTURE Output and Implementing the Evanno Method. Conserv Genet Resour. 2012;4:359–61.

64. Evanno G, Regnaut S, Goudet J. Detecting the number of clusters of individuals using the software STRUCTURE: a simulation study. Mol Ecol. 2005;14:2611–20.

65. Jakobsson M, Rosenberg NA. CLUMPP: A cluster matching and permutation program for dealing with label switching and multimodality in analysis of population structure. Bioinformatics. 2007;23:1801–6.

66. Jombart T, Ahmed I. adegenet 1.3-1: new tools for the analysis of genome-wide SNP data. Bioinformatics. 2011 Nov 1;27(21):3070–1. doi:10.1093/bioinformatics/btr521

67. Jost L. GST and its relatives do not measure differentiation. Mol Ecol. 2008 Sep;17(18):4015–26. doi:10.1111/j.1365-294X.2008.03887.x PubMed PMID: 19238703.

68. Keenan K, Mcginnity P, Cross TF, Crozier WW, Prodöhl PA. DiveRsity: An R package for the estimation and exploration of population genetics parameters and their associated errors. Methods Ecol Evol. 2013;4(8):782–8. doi:10.1111/2041-210X.12067

69. Sundqvist L, Keenan K, Zackrisson M, Prodöhl P, Kleinhans D. Directional genetic differentiation and relative migration. Ecol Evol. 2016 Jun 1;6(11):3461–75. doi:10.1002/ece3.2096

70. Jalón-Rojas I, Wang XH, Fredj E. A 3D numerical model to Track Marine Plastic Debris (TrackMPD): Sensitivity of microplastic trajectories and fates to particle dynamical properties and physical processes. Mar Pollut Bull. 2019 Apr;141:256–72. doi:10.1016/j.marpolbul.2019.02.052

71. Hosegood PJ, Gregg MC, Alford MH. Wind-driven submesoscale subduction at the north Pacific subtropical front. J Geophys Res Oceans. 2013 Oct;118(10):5333–52. doi:10.1002/jgrc.20385

72. Hosegood PJ, Gregg MC, Alford MH. Restratification of the Surface Mixed Layer with Submesoscale Lateral Density Gradients: Diagnosing the Importance of the Horizontal Dimension. J Phys Oceanogr. 2008 Nov 1;38(11):2438–60. doi:10.1175/2008JPO3843.1

73. Hosegood PJ, Nightingale PD, Rees AP, Widdicombe CE, Woodward EMS, Clark DR, et al. Nutrient pumping by submesoscale circulations in the mauritanian upwelling system. Prog Oceanogr. 2017 Dec;159:223–36. doi:10.1016/j.pocean.2017.10.004

74. Wood S, Baums IB, Paris CB, Ridgwell A, Kessler WS, Hendy EJ. El Niño and coral larval dispersal across the eastern Pacific marine barrier. Nat Commun. 2016 Aug 23;7. doi:10.1038/ncomms12571 PubMed PMID: 27550393.

75. Connolly SR, Baird AH. Estimating dispersal potential for marine larvae: dynamic models applied to scleractinian corals. Ecology. 2010.91(12);3572–3583

76. Baird AH, Guest JR, Edwards AJ, Bauman AG, Bouwmeester J, Mera H, et al. An Indo- Pacific coral spawning database. Sci Data. 2021 Dec 1;8(1). doi:10.1038/s41597-020-00793-8 PubMed PMID: 33514754.

77. Koester A, Ford AK, Ferse SCA, Migani V, Bunbury N, Sanchez C, et al. First insights into coral recruit and juvenile abundances at remote Aldabra Atoll, Seychelles. PLoS One. 2021 Dec 1;16. doi:10.1371/journal.pone.0260516 PubMed PMID: 34874982.

78. Oksanen J, Blanchet FG, Kindt R, Legendre P, Minchin PR, O’Hara RB, et al. vegan: Community Ecology Package. Software. 2025. http://CRANR-project.org/package=vegan.

79. Gamoyo M, Obura D, Reason CJC. Estimating Connectivity Through Larval Dispersal in the Western Indian Ocean. J Geophys Res Biogeosci. 2019 Aug 1;124(8):2446–59. doi:10.1029/2019JG005128

80. Treml EA, Roberts JJ, Chao Y, Halpin PN, Possingham HP, Riginos C. Reproductive output and duration of the pelagic larval stage determine seascape-wide connectivity of marine populations. Integr Comp Biol. 2012;52:525–37.

81. Gavrilets S. Fitness Landscapes and the Origin of Species. Princeton University Press. 2004.

82. Templeton AR. The reality and importance of founder speciation in evolution. Bioessays. 2008. 30(5):470–9. doi: 10.1002/bies.20745.

83. Vogt-Vincent NS, Mitarai S, Johnson HL. High-frequency variability dominates potential connectivity between remote coral reefs. Limnol Oceanogr. 2023 Dec 1;68(12):2733–48. doi:10.1002/lno.12455

84. van der Ven RM, Triest L, De Ryck DJR, Mwaura JM, Mohammed MS, Kochzius M. Population genetic structure of the stony coral Acropora tenuis shows high but variable connectivity in East Africa. J Biogeogr. 2016 Mar 1;43(3):510–9. doi:10.1111/jbi.12643

85. van der Ven RM, Ratsimbazafy HA, Kochzius M. Large-scale biogeographic patterns are reflected in the genetic structure of a broadcast spawning stony coral. Coral Reefs. 2022 Jun 1;41(3):611–24. doi:10.1007/s00338-022-02220-3

86. Hata T, Madin JS, Cumbo VR, Denny M, Figueiredo J, Harii S, et al. Coral larvae are poor swimmers and require fine-scale reef structure to settle. Sci Rep. 2017 Dec 1;7(1). doi:10.1038/s41598-017-02402-y PubMed PMID: 28533550.

87. Gilmour JP, Smith LD, Brinkman RM. Biannual spawning, rapid larval development and evidence of self-seeding for scleractinian corals at an isolated system of reefs. Mar Biol. 2009 May 19;156(6):1297–309. doi:10.1007/s00227-009-1171-8

88. Gilmour JP, Smith LD, Heyward AJ, Baird AH, Pratchett MS. Recovery of an Isolated Coral Reef System Following Severe Disturbance. Science. 2013 Apr 5;340(6128):69–71. doi:10.1126/science.1232310

89. Bode M, Leis JM, Mason LB, Williamson DH, Harrison HB, Choukroun S, et al. Successful validation of a larval dispersal model using genetic parentage data. PLoS Biol. 2019 Jul 12;17(7):e3000380. doi:10.1371/journal.pbio.3000380

90. Archer M, Wang J, Klein P, Dibarboure G, Fu LL. Wide-swath satellite altimetry unveils global submesoscale ocean dynamics. Nature. 2025 Apr 17;640(8059):691–6. doi:10.1038/s41586-025-08722-8

91. Grimaldi CM, Lowe RJ, Benthuysen JA, Cuttler MVW, Green RH, Radford B, et al. Hydrodynamic drivers of fine-scale connectivity within a coral reef atoll. Limnol Oceanogr. 2022 Oct 1;67(10):2204–17. doi:10.1002/lno.12198

92. Sola E, Marques da Silva I, Glassom D. Reproductive synchrony in a diverse Acropora assemblage, Vamizi Island, Mozambique. Marine Ecology. 2016 Dec 1;37(6):1373–85. doi:10.1111/maec.12348

93. Mangubhai S, Harrison PL. Asynchronous coral spawning patterns on equatorial reefs in Kenya. Mar Ecol Prog Ser. 2008 May 22;360:85–96. doi:10.3354/meps07385

94. Mangubhai S, Harris A, Graham NAJ. Synchronous daytime spawning of the solitary coral Fungia danai (Fungiidae) in the Chagos Archipelago, central Indian Ocean. Coral Reefs. 2007 Mar;26(1):15. doi:10.1007/s00338-006-0173-y

95. Choukroun S, Stewart OB, Mason LB, Bode M. Larval dispersal predictions are highly sensitive to hydrodynamic modelling choices. Coral Reefs. 2025 Feb 2;44(1):1–13. doi:10.1007/s00338-024-02563-z

96. Meziere Z, Prata K, Lechene M, Ferrari R, Popovic I, Riginos C. Connectivity differs by orders of magnitude among co-distributed corals, affecting spatial scales of eco-evolutionary processes. Sci Adv. 2025 Jul 4;11(27). doi:10.1126/sciadv.adt2066

97. Figueiredo J, Thomas CJ, Deleersnijder E, Lambrechts J, Baird AH, Connolly SR, et al. Global warming decreases connectivity among coral populations. Nat Clim Chang. 2022;12:83–7.

98. Schott FA, Xie SP, McCreary JP. Indian ocean circulation and climate variability. Rev Geophys. 2009;47:1–46.

99. Abram N, Gagan M, Cole J, Hantoro WS, Mudelsee M. Recent intensification of tropical climate variability in the Indian Ocean. Nature Geosci. 2008;849–53.

100. Saura S, Bodin Ö, Fortin MJ. Stepping stones are crucial for species’ long-distance dispersal and range expansion through habitat networks. Journal of Applied Ecology. 2014 Feb;51(1):171–82. doi:10.1111/1365-2664.12179

101. Rodgers EM, Baldanzi S, Collins M, Dowd WW, Feugere L, Mottola G, et al. Protective multi-stressor interactions in the Anthropocene: Key considerations for investigating cross-tolerance in a conservation context. Conserv Physiol. 2025 Jan 1;13(1). doi:10.1093/conphys/coaf052

