## Supplementary figures and tables for "Genomic and oceanographic evidence of reef connectivity across the western and central Indian Ocean": Supplementary material_Coral connectivity_Oct2026.docx

**Supplementary Materials**

**Additional Methods**

*Oceanographic Modelling*

The velocity field output from the Ocean General Circulation Model (OGCM) is used to advect particles from one timestep to the next. Drag is ignored. Advection can be computed using the 1^st^ order Euler scheme, or 2^nd^ or 4^th^ order Runge-Kutta method; in the current analysis we used the 4^th^ order scheme for improved accuracy at the expense of computational time. At first order, the particle position after one timestep, Δt_i_, is **x_n+1_** and obtained from the previous position, **x_n_**, and velocity, **U**, responsible for advection:

$$\boldsymbol{x}_{n+1}=\boldsymbol{x}_{n}+\Delta t_{i}\boldsymbol{U}$$

The 4^th^ order scheme iterates the approximation of particle position based on the lower order estimates.

As the role of submesoscales (71–73), which are dynamical processes with horizontal scales of 1-10 km and that are concentrated around fronts whose circulation accelerates particles within their vicinity and increases diffusivity, cannot be accounted explicitly in the global-scale model we set the horizontal dispersion to 10 m^2^s^-1^. Horizontal dispersion, to some degree at least, allows particle trajectories that to begin with are closely located, start to diverge as they would do when submesoscales are present. Dispersion follows a random-walk model:

$$x_{n+1}=x_{n}+R\sqrt{2K_{h}\Delta t_{i}}$$

where **K_h_** is the horizontal diffusivity, **x_n_** is the original particle position, **R** is the random number with mean zero and a standard deviation, **r=1** and $\boldsymbol{\Delta}$**t**_i_ is the internal timestep as defined above, specified here as 1 day to match that of the OGCM.

In line with previous studies (35,74) we model coral larvae as positively buoyant, otherwise passive particles that drift according to surface currents. Therefore, TrackMPD was run in two-dimensional mode with surface currents only, precluding the effects of sinking. The direct impact of wind on particles is not accounted for in the simulations but the role of wind forcing in generating ocean currents is included in the output from OGCM.

**Tables & Figures**

Table S1. All reef sites and their sample numbers which comprise the six study locations.

| **Locations** | **Reef Sites** | **Number of samples** |
| --- | --- | --- |
| Iles Glorieuses | Grand Glorieuse & Ile du lys | 9 |
| Outer Seychelles Islands | Assumption | 6 |
|  | Aldabra | 18 |
|  | Astove | 3 |
| Inner Seychelles Islands | Praslin Atoll | 5 |
|  | Curieuse | 3 |
|  | Aride | 13 |
|  | Fregate | 10 |
| Diego Garcia | Cannon Point | 6 |
|  | Mid Island | 13 |
|  | Barton Point | 14 |
| Western Chagos | South Brothers & Eagle, Great Chagos Bank | 7 |
|  | Egmont Atoll | 5 |
| Northern Chagos | South Peros Banhos | 24 |
|  | North Peros Banhos | 21 |
|  | West Salomon | 25 |
|  | North Salomon | 26 |
|  | Nelson, Great Chagos Bank | 9 |

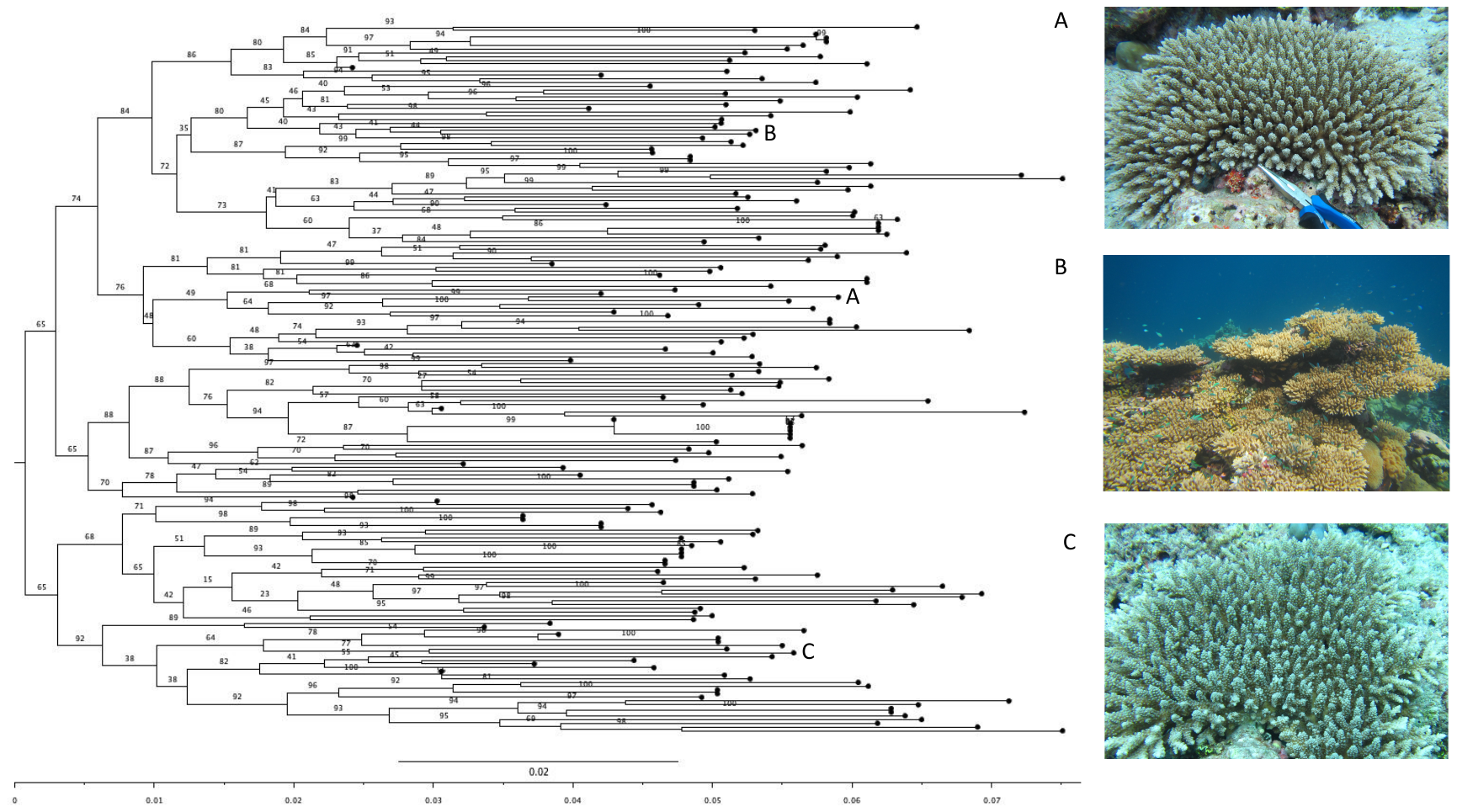

Figure S.1. Midpoint rooted, ML inference tree of final *A.* aff*. tenuis* specimens, with 5000 bootstraps, for downstream pop gen analysis. Photos A-C are examples of *A.* aff. *tenuis* specimens collected, with the corresponding letter marking their location on the tree. Photo B shows a particularly large colony found in the northern Chagos atolls.

Mean LnProb

| K value | LnProb Mean |
| --- | --- |
| 1 | -675280 |
| 2 | -638672 |
| 3 | -631794 |
| 4 | -628777 |
| 5 | -626432 |
| 6 | -627866 |

Figure S.2. Mean linear probability values for K= 1-6

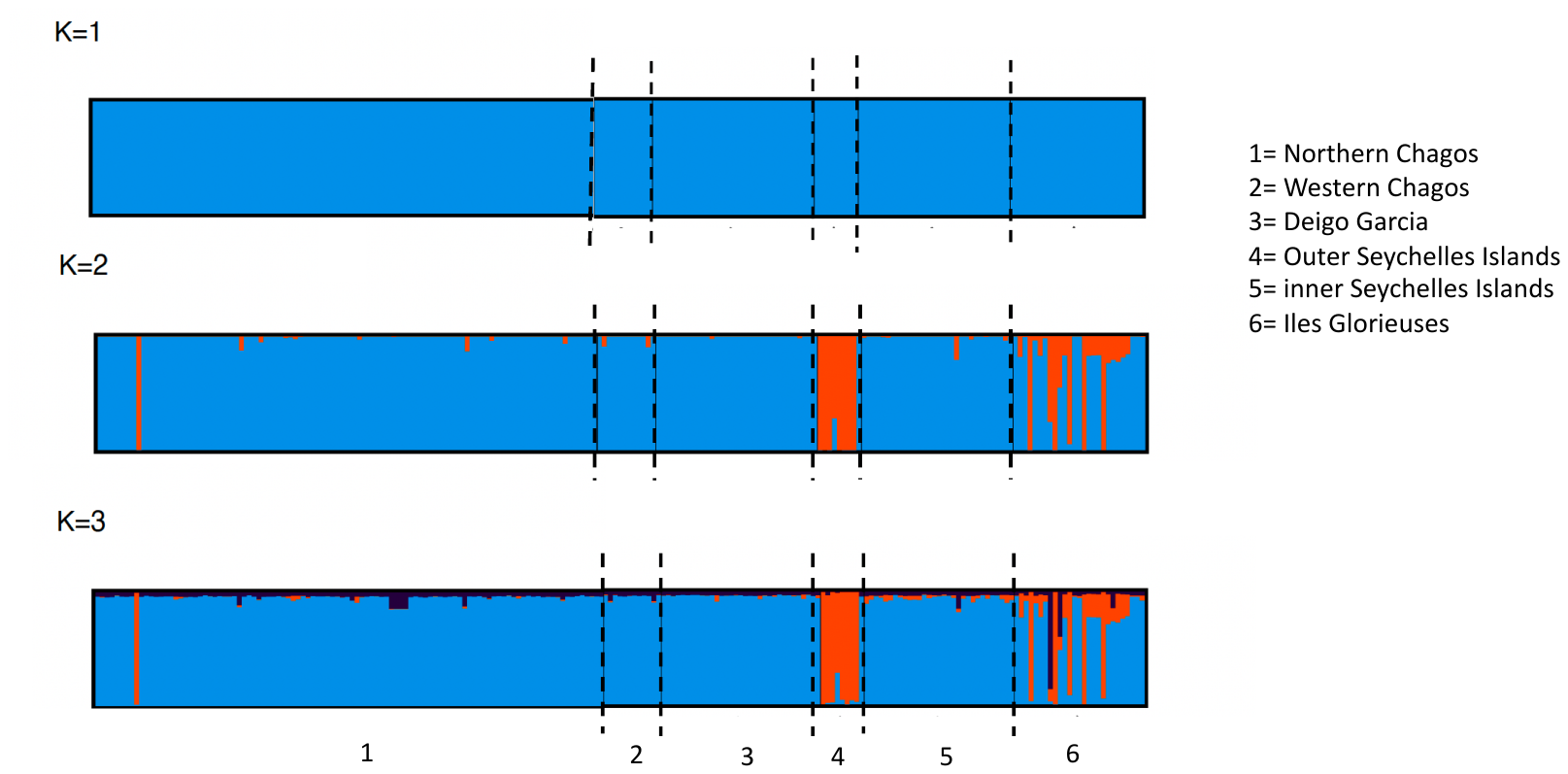

Figure S.3. Barplots of the population structure for K values 1-3.

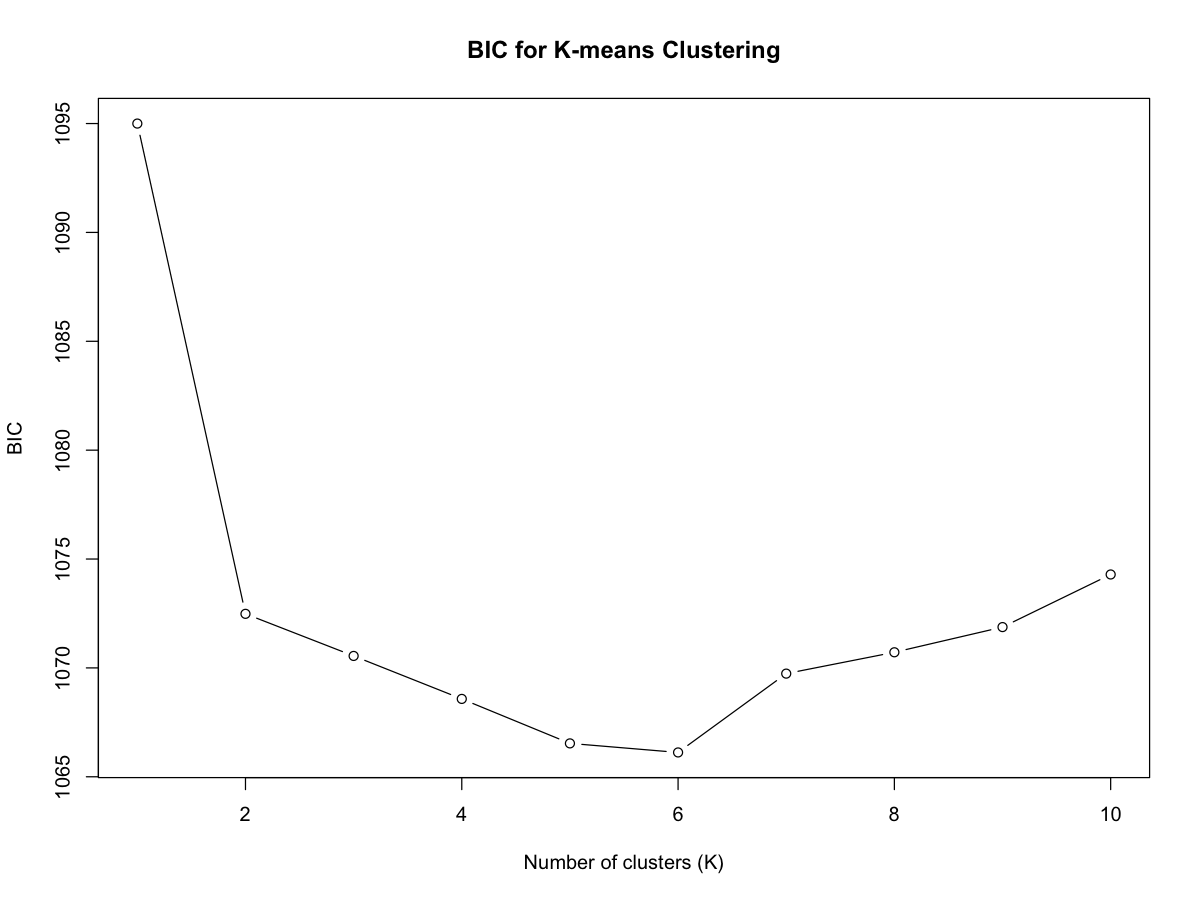

Figure S.4. Graph demonstrating BIC values from the DAPC statistically supporting two clusters across the populations.

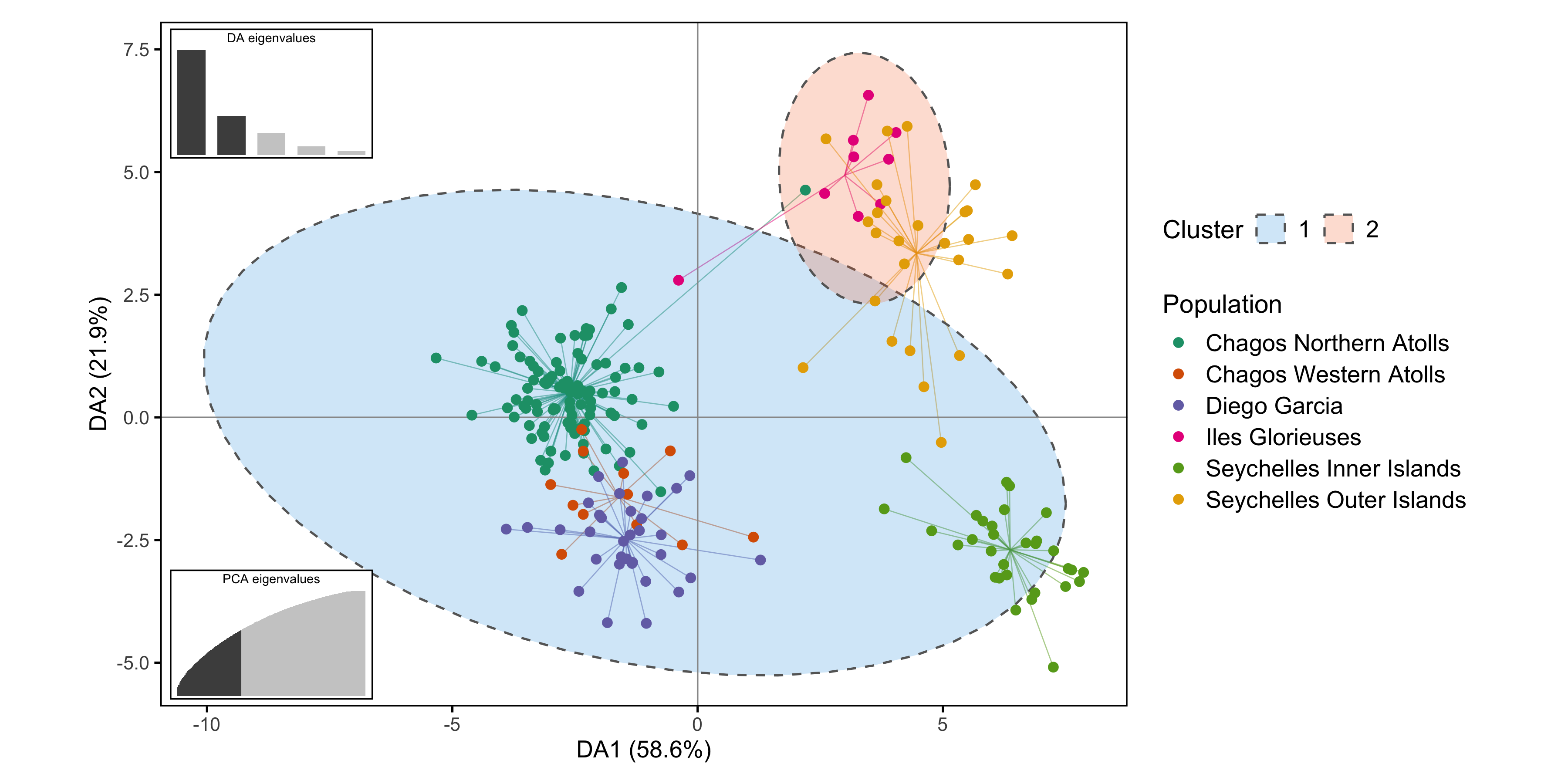

Figure S.5. DAPC showing two major clusters across the six locations statistically supported by BIC. The visual appearance of three clusters reflects a secondary (weaker level) of structure that is real but not statistically compelling enough for BIC to formally split from K=2.

Table S2. F*st* values and Phi-statistics between pairwise populations represented by bold text and normal text respectively. F*st* is the proportion of the total genetic variance contained in a subpopulation relative to the total genetic variance. The greater the F*st* values are, the greater the differences between populations. Phi-statistics is a modified version of Wright's F that refers to the relative contributions of between-population separation to the overall genetic variation in the whole sample. All values can range from 0 to 1.

|  | Northern Chagos | Western Chagos | Diego Garcia | Iles  Glorieuses | Seychelles Inner Islands | Seychelles Outer Islands |
| --- | --- | --- | --- | --- | --- | --- |
| Northern Chagos | Na | **0.01** | **0.01** | **0.04** | **0.01** | **0.02** |
| Western Chagos | 0.00 | Na | **0.01** | **0.11** | **0.02** | **0.02** |
| Diego Garcia | 0.01 | 0.00 | Na | **0.08** | **0.02** | **0.02** |
| Iles Glorieuses | 0.13 | 0.13 | 0.14 | Na | **0.08** | **0.03** |
| Seychelles Inner Islands | 0.02 | 0.01 | 0.01 | 0.13 | Na | **0.02** |
| Seychelles Outer Islands | 0.04 | 0.01 | 0.03 | 0.03 | 0.02 | Na |

Table S3. Migration rates (Jost D) calculated using DivMigrate with 500 bootstraps

| Source Population | Sink: Northern Chagos | Sink: Western Chagos | Sink: Diego Garcia | Sink: Iles Glorieuses | Sink: Inner Seychelles | Sink: Outer Seychelles |
| --- | --- | --- | --- | --- | --- | --- |
| Northern Chagos |  | 0.589 | 0.900 | 0.039 | 0.511 | 0.264* |
| Western Chagos | 0.740 |  | 0.588 | 0.038 | 0.418 | 0.245* |
| Diego Garcia | 1.000* | 0.492 |  | 0.039 | 0.526 | 0.248* |
| Iles Glorieuses | 0.074* | 0.069* | 0.074* |  | 0.077* | 0.147* |
| Inner Seychelles | 0.561 | 0.350 | 0.497* | 0.041 |  | 0.345* |
| Outer Seychelles | 0.218 | 0.166 | 0.195 | 0.061 | 0.244 |  |
| * indicates migration rates that are statistically higher relative to the opposite direction. | | | | | | |

Figure S.6.A & B. Videos tracking larval particle movement in A) negative Indian Ocean Dipole (IOD) phases (29^th^ Oct 2010 – 1^st^ March 2011) and B) positive IOD phases (29^th^ Oct 2019 – 29^th^ Feb 2020), over four months, as an example of how IOD events can change the meanderings of the Southern Equatorial Current (SEC).

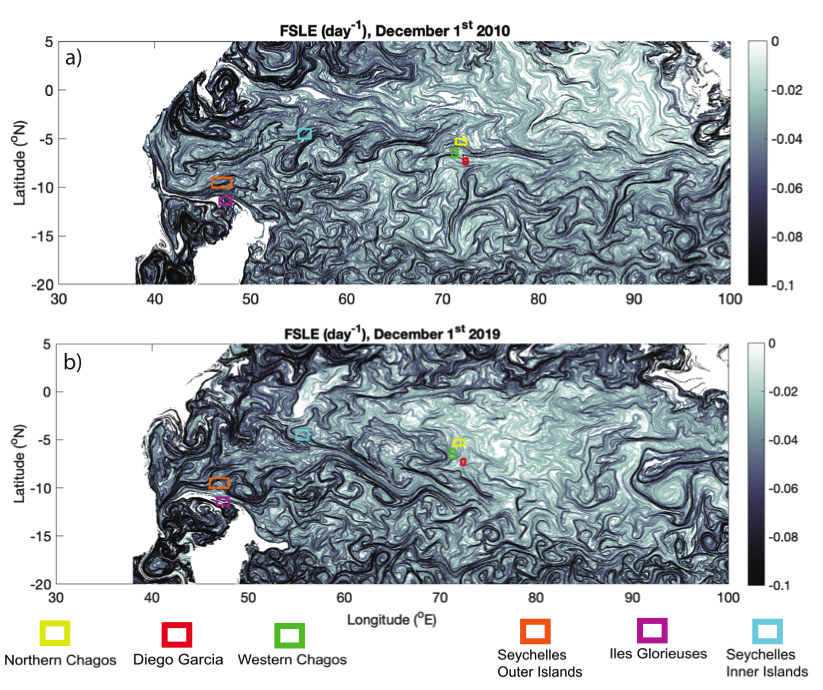

Figure S.7. Provides an example of the sub-mesoscale currents and eddies taking place in the Indian Ocean on any one day which cannot be captured by oceanographic models and hence lead to variation in larvae dispersal
